# High-Throughput Kinase Inhibitor Screening Reveals Roles for Aurora and Nuak Kinases in Neurite Initiation and Dendritic Branching

**DOI:** 10.1101/2020.06.25.162271

**Authors:** Sara M. Blazejewski, Sarah A. Bennison, Xiaonan Liu, Kazuhito Toyo-oka

## Abstract

Kinases are essential regulators of a variety of cellular signaling processes, including neurite formation – a foundational step in neurodevelopment. Aberrant axonal sprouting and failed regeneration of injured axons are associated with conditions like traumatic injury, neurodegenerative disease, and seizures. Investigating the mechanisms underlying neurite formation will allow for identification of potential therapeutics. We used a kinase inhibitor library to screen 493 kinase inhibitors and observed that 45% impacted neuritogenesis in Neuro2a (N-2a) cells. Based on the screening, we further investigated the roles of Aurora kinases A, B, and C and Nuak kinases 1 and 2. The roles of Aurora and Nuak kinases have not been thoroughly studied in the nervous system. Inhibition or overexpression of Aurora and Nuak kinases in primary cortical neurons resulted in various neuromorphological defects, with Aurora A regulating neurite initiation, Aurora B and C regulating neurite initiation and elongation, all Aurora kinases regulating arborization, and all Nuak kinases regulating neurite initiation and elongation and arborization. Our high-throughput screening and analysis of Aurora and Nuak kinases revealed their functions and may contribute to the identification of therapeutics.

## Introduction

Neurite formation is a highly regulated foundational step in neuromorphogenesis that is closely followed by specification of axons and dendrites and establishment of synaptic connections. It is essential that neurite formation occurs with fidelity to allow the subsequent stages of neuromorphogenesis to be properly completed, such as dendritic branching and synapse formation. Overall neuronal morphology can impact connectivity, which affects the amount of inputs received by a neuron and a neuron’s ability to fire action potentials [1, 2]. Neurite formation may be divided into neurite initiation and neurite elongation. Neurite initiation can be associated with a break in the morphological symmetry of a neuron, which drives neuronal polarization. The hallmark of neurite initiation is the extension of actin-rich filopodia and lamellipodia that develop into neurites. These relatively dynamic actin-rich precursor structures are then stabilized via microtubule invasion, which is followed by rapid condensation of the structure to prevent collapse [3]. Once the neurite has been stabilized, neurite elongation will begin as neurites extend and develop growth cones, which are specialized, dynamic regions of actin-rich lamellipodia and filopodia found at the distal ends of all neurites that are critical for forming synaptic contacts and developing neural circuits [4, 5]. Neurite elongation is an extremely dynamic process, which is critical for synaptic plasticity, synaptic pruning, and synapse formation, as well as correct pathfinding [6].

Understanding how neurite formation occurs and is regulated during normal development is necessary for developing therapeutic innovation to either restore neurite formation or trim back aberrant neurite formation. Such treatments would benefit patients suffering from seizures, spinal cord injury, and neurodegenerative disorders that result in axonal injury or degeneration. Neurite formation is controlled by innumerable signaling pathways, with kinases playing an integral role by regulating the phosphorylation status of key target molecules that drive neurite initiation and neurite elongation [7]. Phosphorylation allows for the activation and deactivation of components of a regulatory signaling pathway. Kinases play crucial roles in wide-spread cellular processes including cell cycle regulation, proliferation, metabolism, and apoptosis [8, 9]. There is huge therapeutic potential for kinase inhibitors, as nearly 50 kinase inhibitors have already been approved by the FDA for use [10]. The pharmaceutical industry has recently shifted towards expanding the applicability of pre-approved drugs to reduce drug development time by finding additional uses for drugs that already have established safety profiles. Therefore, a better understanding of the role of kinases in regulating neurite formation may contribute to a speedy transition from basic research to clinical treatment due to the prevalence of FDA approved kinase inhibitors.

The kinase signaling pathways that coordinate neurite formation regulate a variety of processes such as cytoskeletal rearrangement and interaction between actin and microtubules, addition to the plasma membrane, cell adhesion, protein synthesis, and coordination between the cytoskeleton and plasma membrane [11]. These processes need to be precisely regulated to cause the morphological changes that occur during neurite formation. Regulation via protein kinases is advantageous because changes in phosphorylation may occur relatively rapidly, allowing the timing of specific regulatory events to be orchestrated more accurately. This is especially important during neurite formation, since other means of regulation, such as changes in gene expression, would be too slow to adequately regulate some aspects of this process. Since changes in morphology rely heavily on cytoskeletal rearrangements, the regulation of the cytoskeleton has clear importance in neurite formation. Thus, kinases known to regulate microtubule or actin dynamics have been the focus of much investigation.

However, kinases that have traditionally been thought about in different contexts are gaining interest in the neurite formation field. The Aurora and Nuak kinases serve as an example for this. The Aurora kinases are well-characterized as mitotic kinases, while the Nuak kinases are AMPK-related kinases that have an established role in maintaining cellular homeostasis. Recently, Aurora Kinase A has been shown to play a critical role in neuronal migration in the developing cortex [12]. Additionally, Aurora Kinase A promotes neurite elongation in dorsal root ganglion cells and primary cortical neurons by reorganizing microtubules via an atypical protein kinase C (aPKC)-Aurora A-NUDEL1 pathway [13]. Furthermore, Aurora Kinase B overexpression results in extended axonal outgrowth, while pharmacological and genetic impairment of Aurora Kinase B activity caused truncation and aberrant motor axon morphology in zebrafish spinal motor neurons [14]. The role of Aurora Kinase C in neurite formation and neuromorphogenesis is currently unknown. Nuak Kinase 1 and Nuak Kinase 2 have roles in neural tube formation, with *NUAK1* and *NUAK2* double knockout mice exhibiting exencephaly, facial clefting, and spina bifida [15]. Nuak Kinase 1 is also involved in cortical axon arborization *in vivo*, with knockdown or overexpression of NUAK1 drastically reducing or increasing axon branching, respectively [16]. Thus, relatively recent literature implicates Aurora Kinase A in neurite elongation, Aurora Kinase B in regulating axon length, and Nuak Kinase 1 in axon branching.

We conducted a high-throughput screen of 493 kinase inhibitors using the DiscoveryProbeTM Kinase Inhibitor Library (APExBIO), which included inhibitors for both Ser/Thr and Tyr kinases, to identify kinases involved in neurite formation. We found that 45% of kinase inhibitors tested produced a neurite formation phenotype. The Aurora Kinases A, B, and C and Nuak Kinases 1 and 2 were implicated by our screening, prompting further investigation of their roles in neurite formation. Since the Aurora and Nuak kinases have well-established functions in regulating microtubule dynamics during mitosis and current literature indicates that these kinases may also be regulating similar functions during neuritogenesis, there is a need to farther clarify the specific roles of the Aurora and Nuak kinases in neurite initiation and neurite elongation.

To further investigate the Aurora and Nuak kinases and confirm our screening we used a variety of pharmacological inhibitors to target individual kinases, as well as groups of kinases within each family. Our results suggest a crucial role for Aurora Kinase B and Nuak Kinase 1 in neuromorphogenesis, while inhibition of a combination of Aurora Kinase A/B, B/C, and A/B/C, as well as Nuak Kinase1/2 produced significant defects in neuronal morphology. Since our loss-of-function experiments using inhibitors indicate severe defects in neurite formation, we also tested the gain-of-function effects of each kinase on neurite formation. Overexpression of Aurora Kinases A, B, and C in primary cortical neurons revealed that Aurora Kinase A is involved in neurite initiation, while Aurora Kinases A, B, and C have roles in dendritic branching. Overexpression of Nuak Kinases 1 and 2 in primary cortical neurons implicates Nuak Kinase 1 in neurite initiation and dendritic branching. These kinases could potentially serve as therapeutic targets for the treatment of central nervous system injury, seizures, and neurodegenerative disorders.

## Results

### Kinase Inhibitor Screening revealed that Multiple Kinases are involved in Neurite Formation

To identify kinases that are involved in the regulation of neurite formation, we conducted a high-throughput screening by treating Neuro2a (N-2a) mouse neuroblastoma cells, which are induced to extend neurite-like extensions by FBS depletion from culture media, with kinase inhibitors (Fig. 1A). Briefly, N-2a cells were cultured for 1 hour in DMEM (+) media containing kinase inhibitors. After 1 hour, the media was changed to DMEM (-) with inhibitors to induce neurite outgrowth. Each of the 493 kinase inhibitors investigated were tested at concentrations of 0.1, 1, and 10 μM to optimize the dosage. Cells were fixed after 2 hours. Three blinded observers used a scoring system to rank the extent of neurite formation in each condition by assigning each a score from 0-8 (Table 1). Scores were averaged and normalized to controls (see Methods for more detail). We observed that treatment with some inhibitors resulted in either no neurite formation or the formation of short neurites (Fig. 1B). We also observed that some inhibitors caused longer neurites compared to the control, suggesting that the kinases inhibited are negative regulators of neurite formation (Fig. 1B). Of the total 493 kinase inhibitors screened, 222 kinase inhibitors, or 45%, caused a neurite formation phenotype in N-2a cells for at least one of the three inhibitor concentrations tested. The percentage of kinase inhibitors that showed a neurite formation phenotype increased as inhibitor dose increased, and some inhibitors produced phenotypes at multiple or all doses (Fig. 1C). Inhibitors produced either no phenotype, a dose-dependent phenotype, or a consistent phenotype across all doses. Of the 202 kinase inhibitors that showed a dose-dependent phenotype that increased in severity as the dose increased, neurite elongation phenotypes were more common at lower doses (0.1 and 1 μM), while neurite initiation and neurite elongation phenotypes were about equally prevalent at the higher dose (10 μM) (Fig. 1D). Together the tyrosine kinase/adaptors and PI3K/Akt/mTOR signaling pathways accounted for about half of the total 203 kinases that showed a dose-dependent phenotype, with various other pathways accounting for the rest of the dose-dependent group (Fig. 1E). Of the 45% of kinase inhibitors that caused a neurite formation phenotype for at least one inhibitor concentration, 19.37% inhibited neurite initiation and 45.50% inhibited neurite elongation in a dose dependent manner, 26.13% inhibited both neurite elongation and initiation in a dose dependent manner, 2.70% inhibited neurite initiation at all doses, 2.25% inhibited neurite elongation at all doses, and 4.05% acted as negative regulators to increase neurite formation (Fig. 1F) (Table 2). The screening also suggested prevalent roles in neurite formation for certain signaling pathways across all groups, including dose-dependent inhibitors, inhibitors of negative regulators, and inhibitors with consistent phenotypes at multiple doses, with the tyrosine kinase/adaptors, JAK/STAT, membrane transporter/ion channel, and chromatin/epigenetics signaling pathways being the top pathways shown to involve kinases that regulate neurite formation overall (Fig. 1G) (Suppl. Table 1).

**Table 1:**
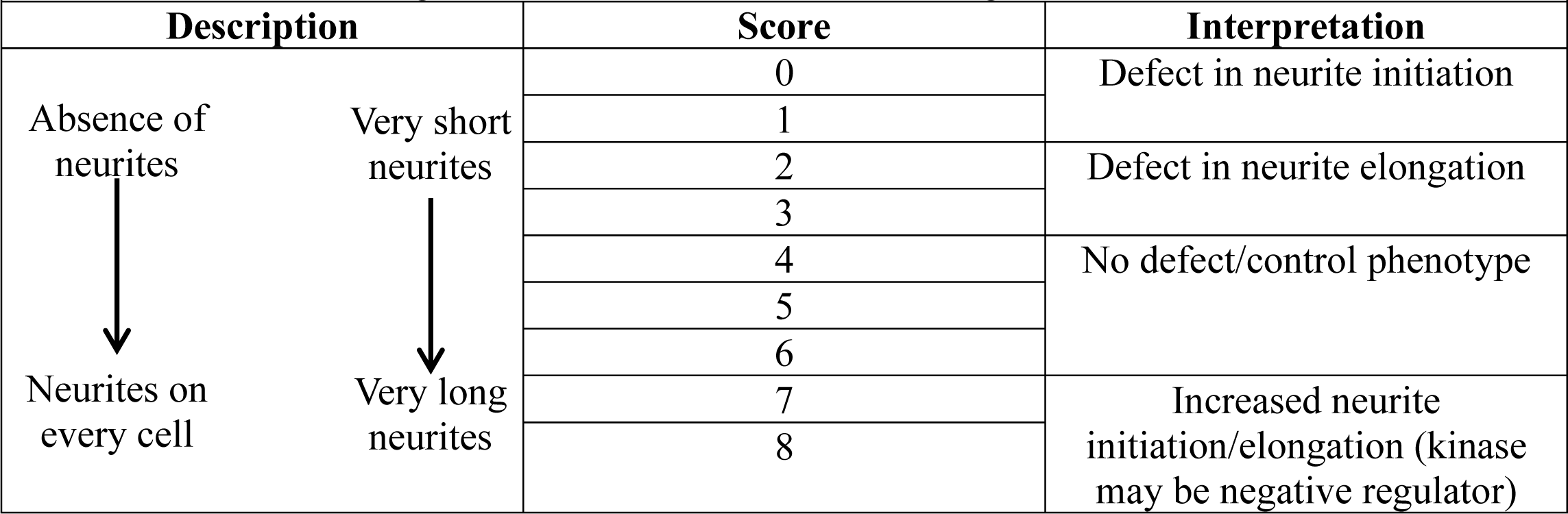
Scale for scoring number of neurites and neurite length in N-2a cells. Scale used by three blind observers for scoring number of neurites from soma and neurite length in N2a cells. The average score for each condition was calculated and used to determine if each kinase inhibitor had an effect on the extent of neurite initiation or elongation.

**Table 2:** Kinase inhibitors that are negative regulators or that have a dose-independent phenotype. Kinase inhibitors that were determined to be negative regulators of neurite formation at 10 μM and regulators of neurite initiation or elongation at all doses (0.1, 1, 10 μM).

| **Table 2:** Kinase inhibitors that are negative regulators or that have a dose-independent phenotype |  |  |
| --- | --- | --- |
| **Kinase inhibitor** | **Pathway** | **Target Kinase** |
| **Negative Regulators (10 μM)** |  |  |
| SNS-032 (BMS-387032) | Cell Cycle/Checkpoint | Cyclin-Dependent Kinases |
| D4476 | Stem Cell | CK1 |
| BX795 | PI3K/Akt/mTOR Signaling | Akt |
| R406 | Tyrosine Kinase/Adaptors | Spleen Tyrosine Kinase (Syk) |
| GNF 5 | TGF-β / Smad Signaling | Bcr-Abl |
| CCT128930 | PI3K/Akt/mTOR Signaling | Akt |
| LEE011 | Cell Cycle/Checkpoint | Cyclin-Dependent Kinases |
| Icotinib | JAK/STAT Signaling | EGFR |
| NSC 23766 | Cell Cycle/Checkpoint | Rho |
| **Regulators of Neurite Initiation (all doses)** |  |  |
| WAY-600 | PI3K/Akt/mTOR Signaling | mTOR |
| TCS JNK 5a | MAPK Signaling | JNK |
| KX2-391 | Tyrosine Kinase | Src |
| KX2-391 dihydrochloride | Tyrosine Kinase/Adaptors | Src |
| Tivantinib (ARQ 197) | Tyrosine Kinase/Adaptors | c-MET |
| Nocodazole | Ubiquitination | Autophagy |
| **Regulators of Neurite Elongation (all doses)** |  |  |
| PP 1 | Tyrosine Kinase/Adaptors | Src |
| ZCL278 | Cell Cycle/Checkpoint | Cdc42 |
| PH-797804 | MAPK Signaling | p38 |
| ML347 | TGF-β / Smad Signaling | TGF-βR1(ALK5) |
| Torin 2 | PI3K/Akt/mTOR Signaling | mTOR |

**Figure 1.**
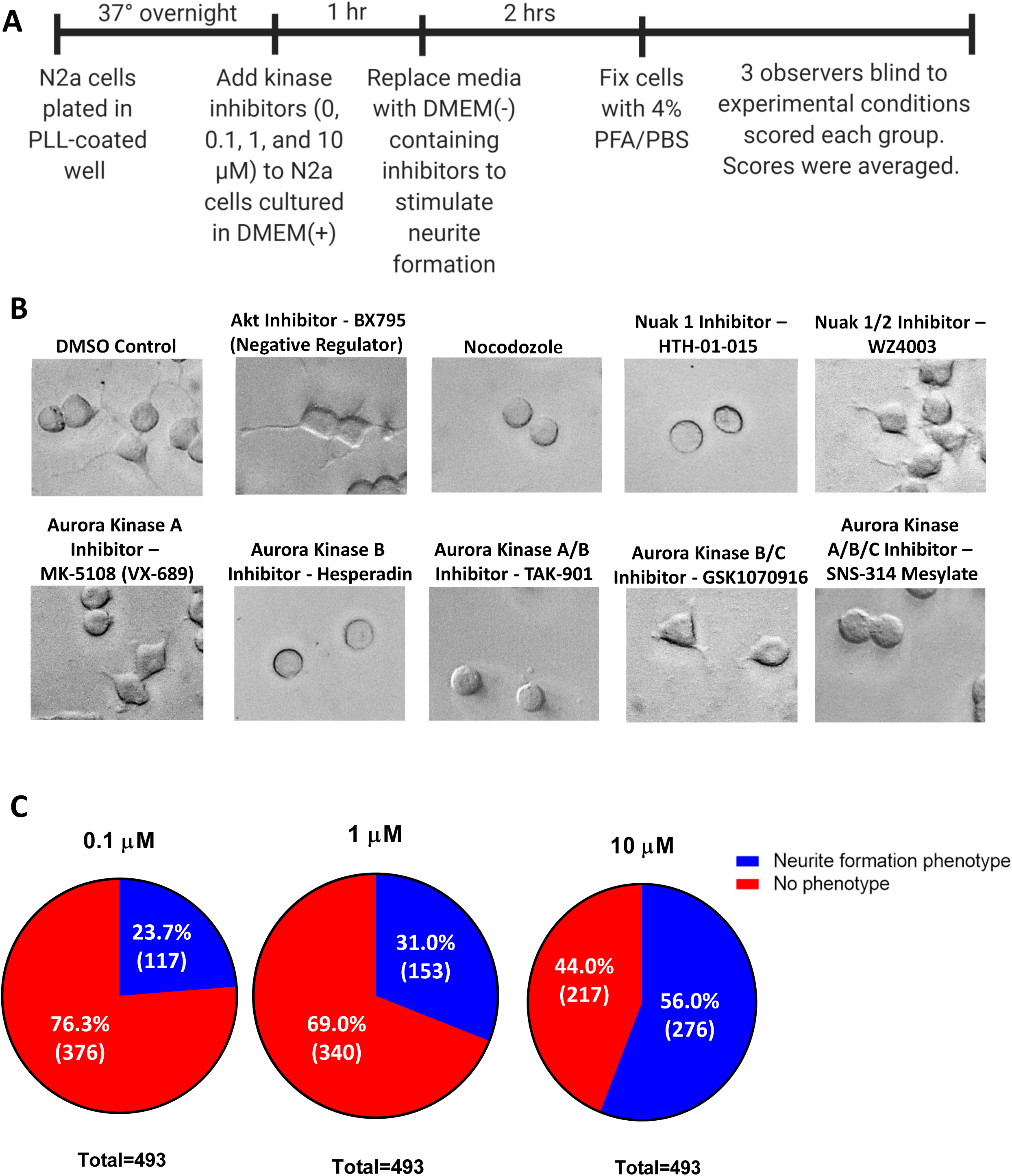
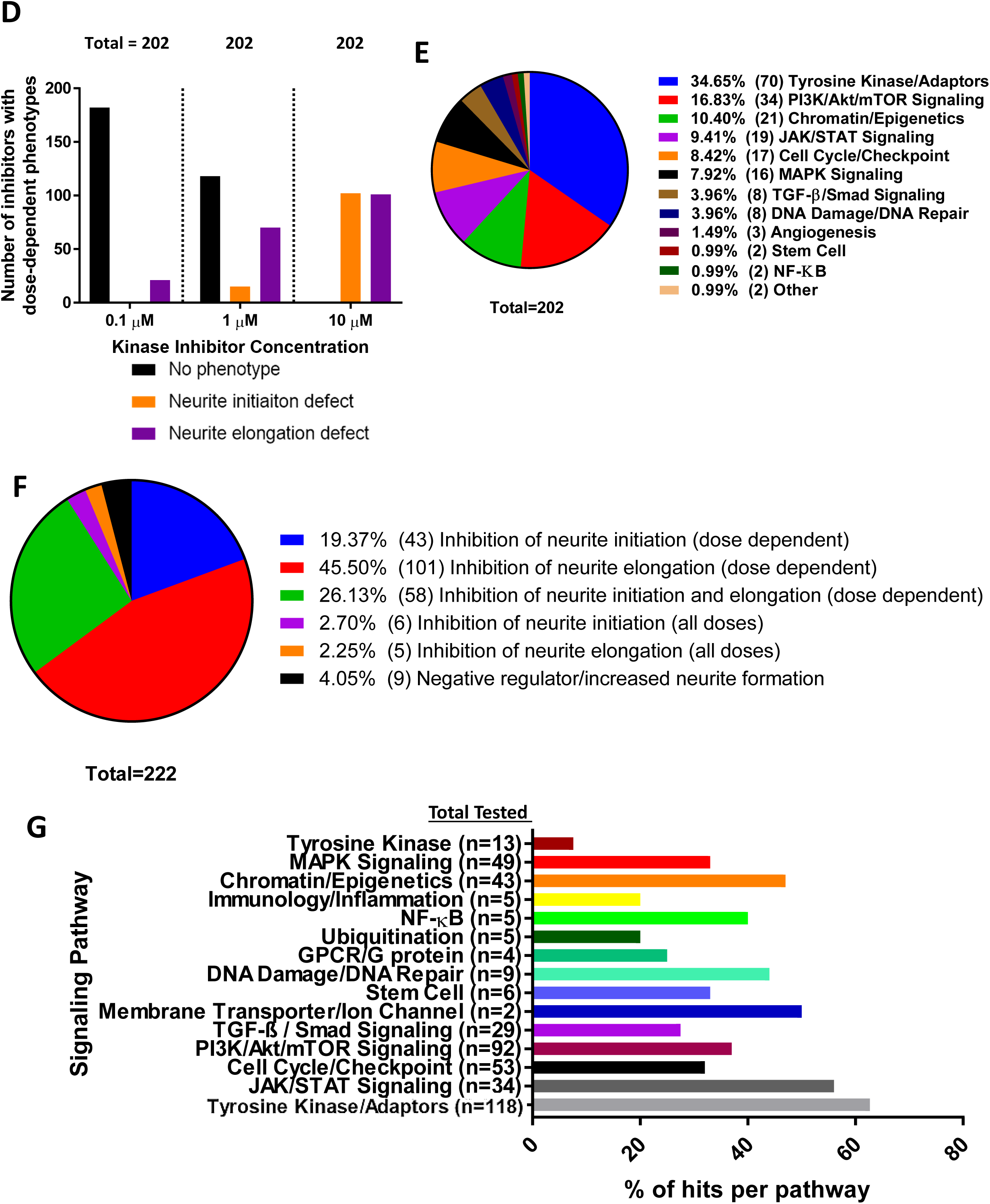
Kinase Inhibitor Screening Revealed Neurite Formation Phenotype for 222 Kinase Inhibitors. A) Timeline describing the experimental design of the kinase inhibitor screening. B) N-2a cells treated with 5 μM DMSO, Akt inhibitor - BX795, Nocodozole, Nuak 1 inhibitor - HTH-01-015, Nuak 1/2 inhibitor - WZ4003, Aurora Kinase A inhibitor - MK-5108 (VX-689), Aurora Kinase B inhibitor - Hesperadin, Aurora Kinase A/B inhibitor - TAK-901, Aurora Kinase B/C inhibitor - GSK1070916, Aurora Kinase A/B/C inhibitor - SNS-314 Mesylate. C) Pie charts showing the percentage of groups that showed a neurite formation phenotype at each inhibitor dose. D) Graph showing the number of dose-dependent inhibitors that had no phenotype, neurite initiation defect, or neurite elongation defect for each inhibitor dose group. E) Pie chart showing the percentage of each signaling pathway that makes up the dose-dependent inhibitors group. F) Summary of kinase inhibitor screening results shows the percentage of kinase inhibitors that resulted in each neurite formation phenotype. Percentages were taken out of the 45% of all inhibitors tested that had an effect on neurite formation, not the total number of inhibitors tested. G) Summary of signaling pathways that kinases implicated in the regulation of neurite formation were most commonly found in, which displays the number of total kinase inhibitors tested on kinases in each signaling pathway and the percentage of those kinase inhibitors that produced a neurite formation phenotype for each. Data from dose-dependent inhibitors, inhibitors of negative regulators, and inhibitors with consistent phenotypes across all doses are represented.

The mean neurite formation scores were used to identify kinases involved in regulating the two stages of neurite formation: neurite initiation and neurite elongation. We considered the effect that kinase inhibitors had on both the number and length of neurites. No or primitive neurite formation reflects a defect in neurite initiation, while decreased neurite length is considered to be a defect in neurite elongation (Table 1). Based on the scores, we selected the kinases in which treatment with inhibitors targeting that kinase resulted in severe defects in neurite formation to investigate further. In some cases, multiple inhibitors in the library targeted the same kinase. The reported mean scores represent the average of all inhibitors that target each kinase.

Our kinase inhibitor screening in N-2a cells revealed neurite formation defects when the Aurora Kinases A, B, and/or C and Nuak Kinases 1 and/or 2 were inhibited at multiple doses (0.1, 1, 10 μM), suggesting a potential role for these kinases in neurite formation (Table 3). To inhibit Aurora Kinases, we used kinase inhibitors targeting either Aurora Kinase A (MK-5108 (VX-689)), B (Hesperadin), A/B (TAK-901), B/C (GSK1070916), or A/B/C (SNS-314 Mesylate) (Fig. 1B). There were 25 kinase inhibitors that target Aurora kinases included in the inhibitor library. The mentioned kinase inhibitors were chosen to display representative images of the screening (Fig. 1B). MK-5108 (VX-689) is a novel, potent and selective inhibitor of Aurora A kinase that competitively binds to its ATP binding site [17]. Hesperadin is an ATP-competitive small molecule inhibitor of Aurora B kinase [18, 19]. TAK-901 is derived from the novel azacarboline kinase hinge binder and has been shown to inhibit Aurora kinase A and B [20]. GSK1070916 is a potent, selective, and reversible ATP-competitive inhibitor of Aurora kinase B and C [21]. SNS-314 Mesylate is an ATP-competitive and selective inhibitor of Aurora kinase A, B, and C [22]. Inhibition of Aurora Kinase A resulted in neurite elongation defects, while inhibition of Aurora Kinase B resulted in neurite initiation defects. However, when both Aurora Kinase A and Aurora Kinase B were inhibited, neurite initiation defects were observed. Additionally, combined inhibition of Aurora Kinase B and Aurora Kinase C led to neurite elongation defects, while inhibition of all Aurora kinases A, B, and C caused neurite initiation defects. This suggests that the specific member or combination of members that are inhibited in the Aurora kinase family impacts the extent that neurite initiation and/or neurite elongation is affected. To inhibit Nuak kinases, we used kinase inhibitors targeting either Nuak Kinase 1 (HTH-01-015) or 1/2 (WZ4003). HTH-01-015 shows extreme selectivity for Nuak 1 and inhibits its phosphorylation, WZ4003 is a potent and selective inhibitor of Nuak kinase 1 and 2 [23]. When Nuak kinase 1 was inhibited, defects in neurite initiation were observed. Neurite elongation deficits were caused upon inhibition of both Nuak kinases 1 and 2. Thus, inhibition of both Nuak kinase 1 and Nuak kinase 2 results in a more severe phenotype.

**Table 3:** Mean neurite formation defect scores for inhibitors targeting Aurora and Nuak kinases across all doses (0.1, 1, 10 μM) Mean neurite formation defect scores for kinase inhibitors targeting the Aurora and Nuak kinases reveal possible defects in neurite initiation and/or elongation.

| **Kinase Inhibitor Targets** | **Average Score** | **Possible Defect** |
| --- | --- | --- |
| DMSO Control | 5.04 | N/A |
| Aurora Kinase A | 3.69 | Neurite elongation |
| Aurora Kinase B | 1.22 | Neurite initiation |
| Aurora Kinase A/B | 1.18 | Neurite initiation |
| Aurora Kinase B/C | 3.74 | Neurite elongation |
| Aurora Kinase A/B/C | 1.46 | Neurite initiation |
| Nuak 1 | 1.09 | Neurite initiation |
| Nuak 1/2 | 3.43 | Neurite elongation |

### Inhibition of Aurora Kinases in Primary Cortical Neurons Suggests Roles for Aurora B and C in Neurite Initiation, Elongation and Arborization

To further investigate the roles of the Aurora Kinases and confirm the results of our N-2a cell screening, neurons expressing YFP were treated with kinase inhibitors targeting either Aurora Kinase A (MK-5108 (VX-689)), B (Hesperadin), A/B (TAK-901), B/C (GSK1070916), or A/B/C (SNS-314 Mesylate) (Fig. 2A). We considered the longest neurite the neurite most likely to become the axon, with the remaining neurites likely to become dendrites. Neurons treated with an Aurora Kinase B inhibitor (10.6 ± 0.95), Aurora Kinase B/C inhibitor (7.7 ± 0.86), or Aurora Kinase A/B/C inhibitor (15.5 ± 1.21) had significantly shorter neurites likely to become dendrites as compared to the DMSO treated control (21.0 ± 1.29) (Fig. 2B). Length of neurites likely to become axons was similarly affected, as neurons treated with an Aurora Kinase B inhibitor (25.7 ± 4.08), Aurora Kinase B/C inhibitor (19.6 ± 1.36), or Aurora Kinase A/B/C inhibitor (23.4 ± 1.73) had significantly shorter neurites likely to become axons as compared to the DMSO treated control (54.1 ± 5.33) (Fig. 2C). Additionally, the number of neurites was significantly decreased in neurons treated with an Aurora Kinase A/B inhibitor (3.68 ± 0.47), Aurora Kinase B/C inhibitor (3.25 ± 0.32), and Aurora Kinase A/B/C inhibitor (2.95 ± 0.28) as compared to the DMSO treated controls (5.60 ± 0.39) (Fig. 2D). While most experimental groups showed a notable decrease in arborization, neurons treated with the Aurora Kinase A inhibitor showed the mildest phenotype with a Sholl profile like DMSO treated control neurons (Fig. 2E). These data suggest that Aurora Kinase B and C are the major isoforms involved in multiple stages of neurite morphogenesis in primary cortical neurons.

**Figure 2.**
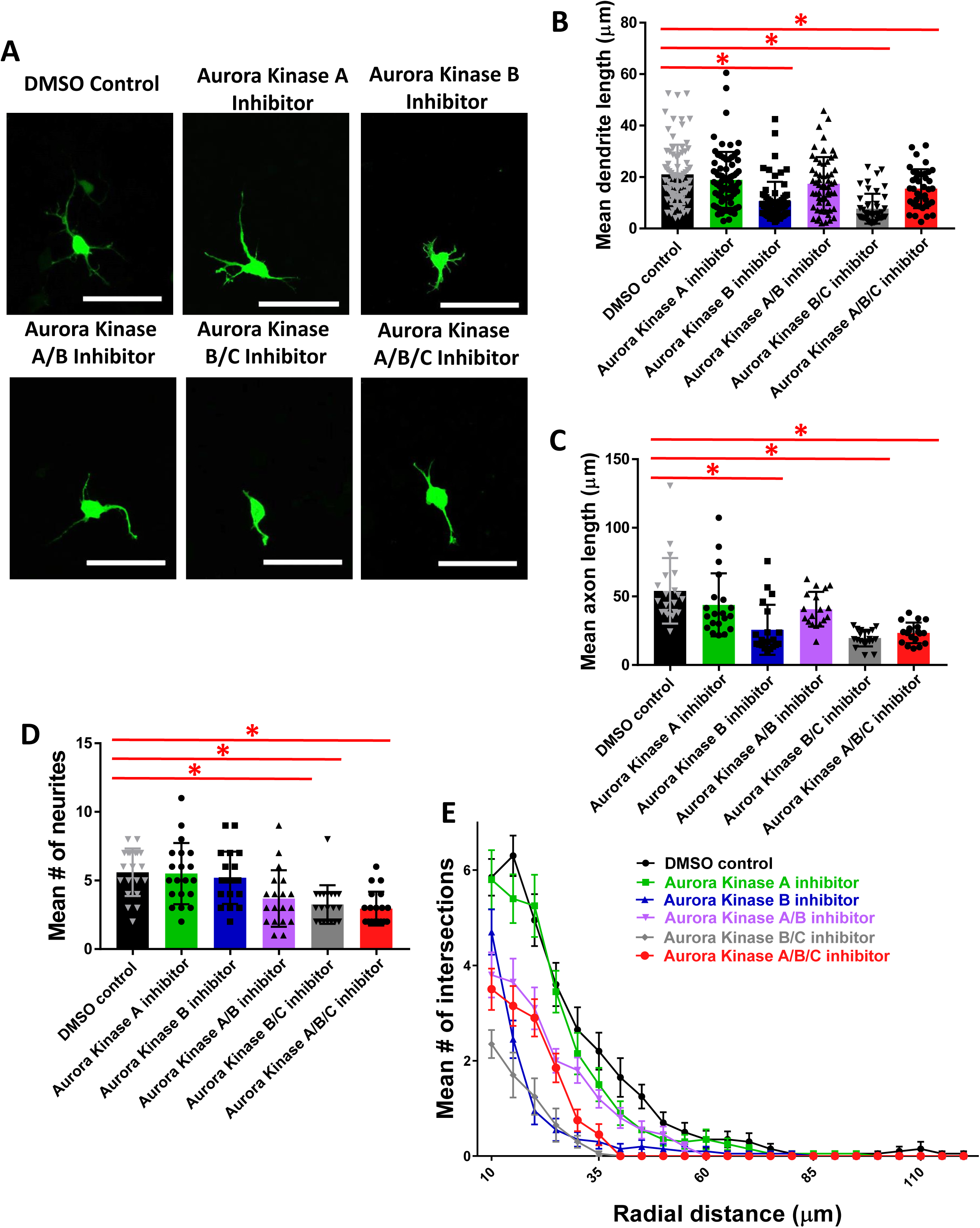
Inhibition of Aurora Kinases A, B, A/B, B/C, and A/B/C in Primary Cortical Neurons Suggests Roles for Aurora B and C in Neurite Initiation/Elongation and Arborization. A) Primary cortical mouse neurons transfected with YFP and treated with Aurora Kinase inhibitors, scale bars=50 μm. B) One-way ANOVA showed significant difference in length of neurites likely to become dendrites (F(5,354)=16.941, *p*<0.0005). There was a significant decrease in neurons treated with Aurora B inhibitor (10.6 ± 0.95, n=64, *p*<0.0005), Aurora B/C inhibitor (7.7 ± 0.86, n=46, *p*<0.0005), or Aurora A/B/C inhibitor (15.5 ± 1.21, n=39, *p*=0.49) compared to DMSO control (21.0 ± 1.29, n=80). There was no significant difference between neurons treated with Aurora A inhibitor (18.9 ± 1.26, n=74, *p*=1.000) or Aurora A/B inhibitor (17.4 ± 1.38, n=57, *p*=0.437) compared to DMSO control. C) One-way ANOVA showed significant difference in length of neurites likely to become axons (F(5,111) = 12.946, *p*<0.0005). There was a significant decrease in neurons treated Aurora B inhibitor (25.7 ± 4.08, n=20, *p*<0.0005), Aurora B/C inhibitor (19.6 ± 1.36, n=20, *p*<0.0005), or Aurora A/B/C inhibitor (23.4 ± 1.73, n=19, *p*<0.0005) compared to DMSO treated control (54.1 ± 5.33, n=20). There was no significant difference between neurons treated with Aurora A inhibitor (43.8 ± 5.13, n=20, *p*=0.860) or Aurora A/B inhibitor (40.7 ± 2.98, n=18, *p*=0.247) compared to DMSO control. D) One-way ANOVA showed significant difference in mean number of neurites (F(5,113) = 8.941, *p*<0.0005). There was a significant decrease in neurons treated with Aurora A/B inhibitor (3.68 ± 0.47, n=19, *p*=0.017), Aurora B/C inhibitor (3.25 ± 0.32, *p*=0.001), and Aurora A/B/C inhibitor (2.95 ± 0.28, *p*<0.0005) compared to the DMSO treated controls (5.60 ± 0.39). There was no significant difference between neurons treated with Aurora A inhibitor (5.50 ± 0.50, *p*=1.000) or Aurora B inhibitor (5.20 ± 0.43, *p*=1.000) and DMSO control. For all groups, n=20 unless otherwise noted. E) Sholl analysis shows dramatic decrease in branching after treatment with Aurora Kinase B, A/B, B/C, and A/B/C inhibitors and a mild decrease in branching after Aurora Kinase A inhibitor treatment.

### Inhibition of Nuak Kinases 1 or 1/2 in Primary Cortical Neurons Suggests Roles for Nuak 1 and 2 in Neurite Initiation and Elongation and Arborization

To further investigate the roles of the Nuak Kinases and confirm the results of our N-2a cell screening, neurons expressing YFP were treated with kinase inhibitors targeting either Nuak Kinase 1 (HTH-01-015) or 1/2 (WZ4003) (Fig. 3A). Neurons treated with a Nuak Kinase 1 inhibitor (4.7 ± 0.54) or Nuak Kinase 1/2 inhibitor (6.48 ± 0.90) had significantly shorter neurites likely to become dendrites as compared to the DMSO treated control (23.1 ± 1.67) (Fig. 3B). Length of neurites likely to be axons was similarly affected, as neurons treated with a Nuak Kinase 1 inhibitor (10.3 ± 1.17) or Nuak Kinase 1/2 inhibitor (15.6 ± 2.19) had significantly shorter neurites likely to be axons as compared to the DMSO treated control (56.2 ± 5.64) (Fig. 3C). Additionally, the number of neurites was significantly decreased in neurons treated with a Nuak Kinase 1 inhibitor (2.65 ± 0.30) or Nuak Kinase 1/2 inhibitor (2.05 ± 0.33) as compared to the DMSO treated control (5.15 ± 0.33) (Fig. 3D). Neurons treated with either the Nuak Kinase 1 or Nuak Kinase 1/2 inhibitor showed a dramatic decrease in arborization (Fig. 3E). These data strongly indicate the importance of the Nuak kinases in neurite morphogenesis in primary cortical neurons.

**Figure 3.**
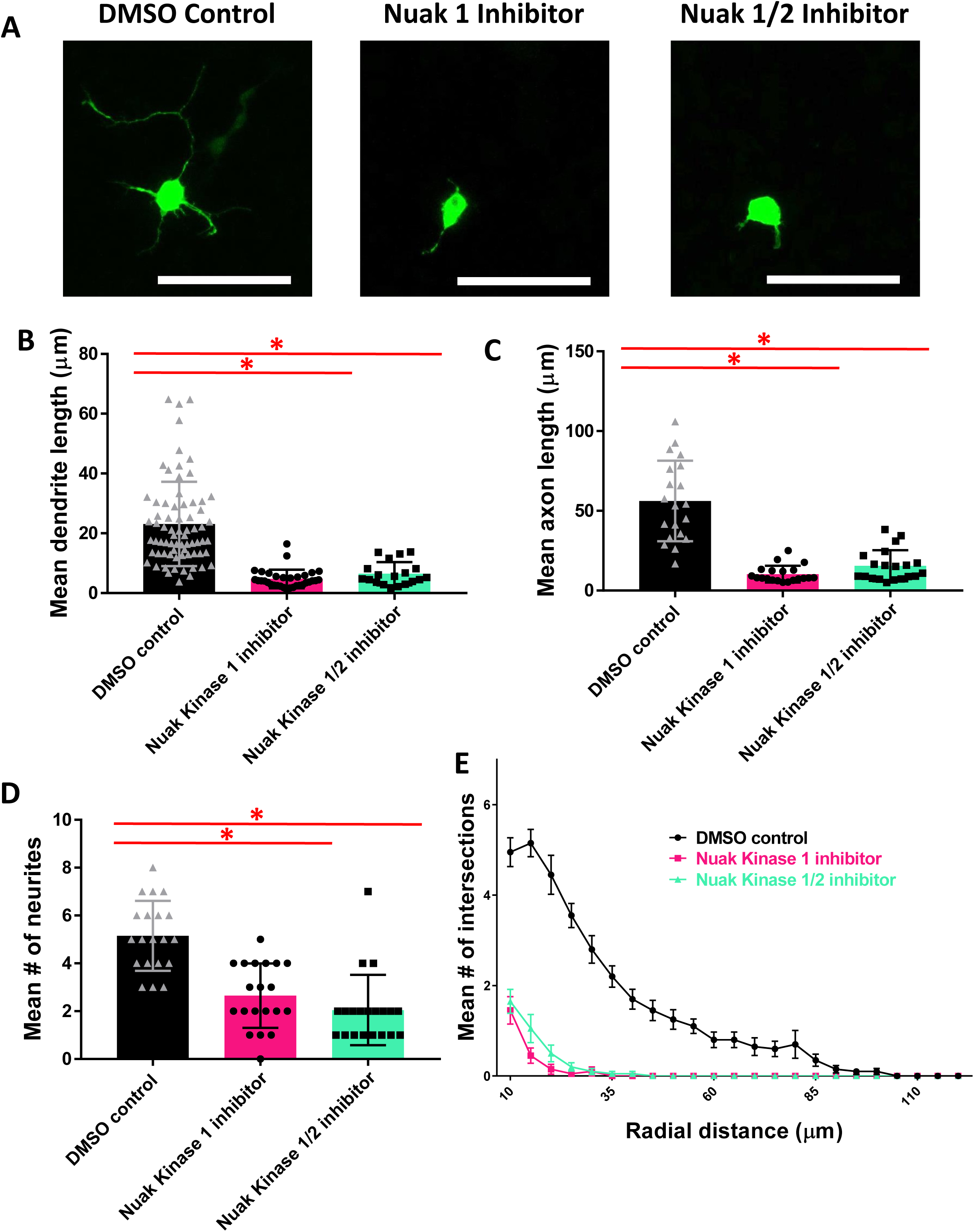
Inhibition of Nuak Kinases 1 and 1/2 in Primary Cortical Neurons Suggests Roles for Nuak 1 and 2 in Neurite Initiation and Elongation and Arborization. A) Primary cortical mouse neurons transfected with YFP and treated with Nuak Kinase inhibitors, scale bars = 50 μm. B) One-way ANOVA determined there was a statistically significant difference in length of neurites likely to become dendrites (F(2, 120) = 39.341, *p*<0.0005). There was a statistically significant decrease in neurons treated with Nuak Kinase 1 inhibitor (4.7 ± 0.54, n=33, *p*<0.0005) and Nuak Kinase 1/2 inhibitor (6.48 ± 0.90, n=19, *p*<0.0005) as compared to the DMSO treated control (23.12 ± 1.67, n=71). C) One-way ANOVA determined there was a statistically significant difference in length of neurites likely to become axons (F(2, 57) = 49.832, *p*<0.0005). There was a statistically significant decrease in neurons treated with Nuak Kinase 1 inhibitor (10.3 ± 1.17, *p*<0.0005) and Nuak Kinase 1/2 inhibitor (15.6 ± 2.19, *p*<0.0005) as compared to the DMSO treated control (56.2 ± 5.64). For all groups, n=20. D) One-way ANOVA determined there was a statistically significant difference in mean number of neurites (F(2, 57) = 26.556, *p*<0.0005). There was a statistically significant decrease in neurons treated with Nuak Kinase 1 inhibitor (2.65 ± 0.30, *p*<0.0005) and Nuak Kinase 1/2 inhibitor (2.05 ± 0.33, *p*<0.0005) as compared to the DMSO treated control (5.15 ± 0.33). For all groups, n=20. E) Sholl analysis shows dramatic decrease in branching for neurons treated with Nuak Kinase 1 and 1/2 inhibitors.

### Overexpression of Aurora Kinases A, B, or C in Primary Cortical Neurons Suggests Role for Aurora Kinase A in Neurite Initiation and for Aurora Kinase A, B, and C in Dendritic Branching

Since our experiments using inhibitors indicate that Aurora kinases are a positive regulator of multiple steps of neurite morphogenesis, we investigated the effects of overexpression of the Aurora kinases on neurite formation to assess the therapeutic potential of activating specific isoforms to increase neurite initiation and/or elongation. Mouse primary cortical neurons were used to overexpress the Aurora kinases A, B, or C (Fig. 4A). The length of neurites likely to become dendrites and neurites likely to become the axon, as well as number of neurites and arborization were analyzed, however no significant differences were found in the length of neurites likely to become dendrites and the axon (Fig. 4 B and C). Interestingly, neurons overexpressing Aurora Kinase A (5.4 ± 0.418) had significantly more neurites as compared to controls (3.6 ± 0.170) (Fig. 4D). The Aurora Kinases A, B, and C all showed increased dendritic branching proximally and decreased dendritic branching distally (Fig. 4E). These data suggest that Aurora kinase A is involved in neurite initiation and that all Aurora kinases are involved in dendritic branching, but have different roles in proximal and distal regions of dendrites.

**Figure 4.**
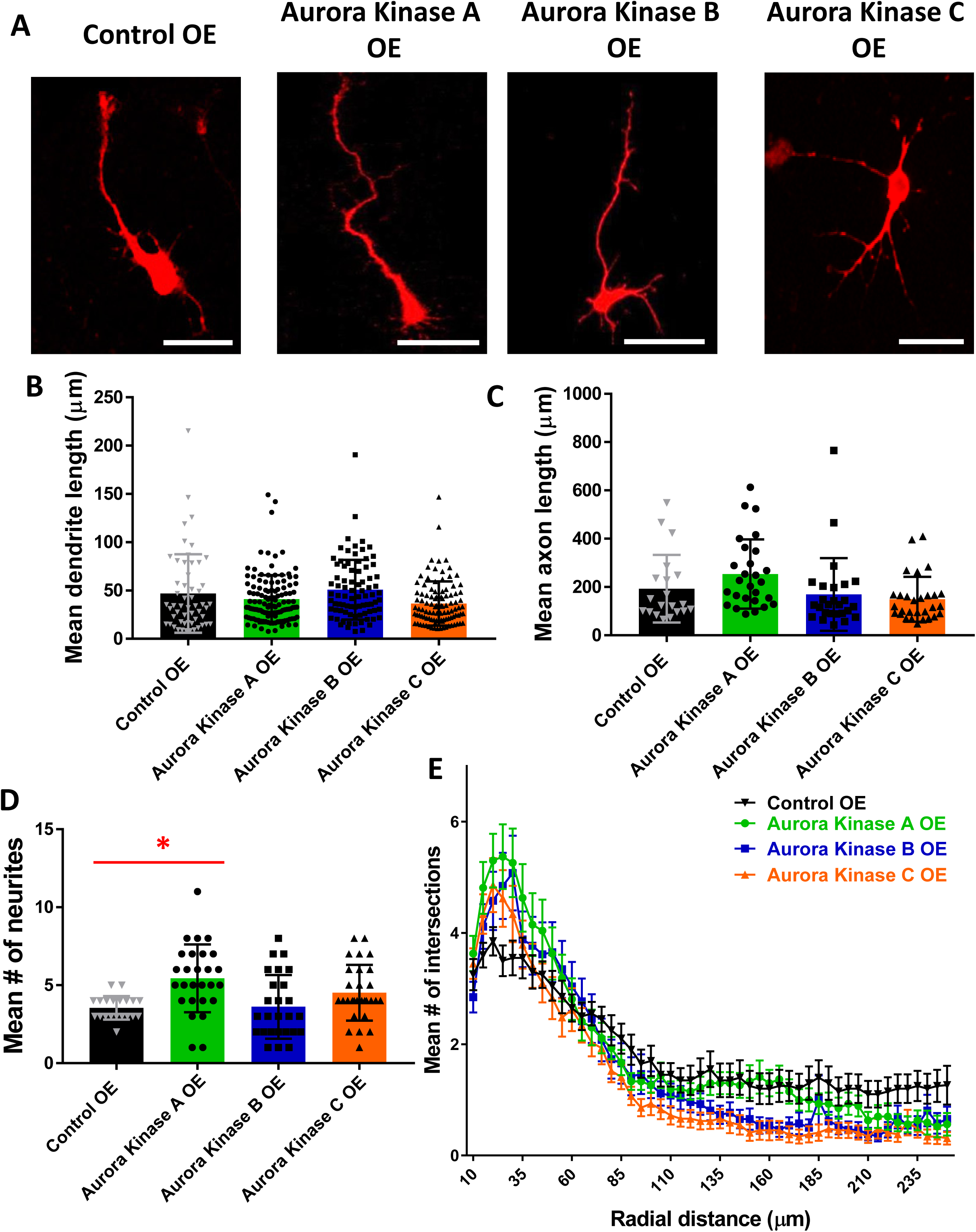
Overexpression of Aurora Kinases A, B, and C in Primary Cortical Neurons Suggests Role for Aurora Kinase A in Neurite Initiation and for Aurora Kinase A, B, and C in Dendritic Branching. A) Primary cortical mouse neurons transfected with mRFP at 48 hours after re-plating, scale bars=50 μm. B) One-way ANOVA determined there was a statistically significant difference in length of neurites likely to become dendrites (F(3,354) = 4.256, *p*=0.006). However, there was no statistically significant difference between Aurora Kinase A OE (41.06 ± 2.28, n=120, *p*=1.000), Aurora Kinase B OE (50.84 ± 3.41, n=82, *p*=1.000), and Aurora Kinase C OE (36.56 ± 2.27, n=102, *p*=0.197) compared to control OE (46.92 ± 5.54, n=54). C) One-way ANOVA determined there was a statistically significant difference in length of neurites likely to become axons (F(3,98) = 3.209, *p*=0.026). However, there was no statistically significant difference between Aurora Kinase A OE (253.54 ± 27.58, n=27, *p*=0.745), Aurora Kinase B OE (169.50 ± 29.48, n=26, *p*=1.000), and Aurora Kinase C OE (149.26 ± 17.27, n=29, *p*=1.000) compared to control OE (193.05 ± 31.27, n=20). D) One-way ANOVA determined there was a statistically significant difference (F(3,98) = 5.991, *p*=0.001). Bonferroni post hoc test revealed that mean number of neurites was statistically significantly greater for Aurora Kinase A OE (5.4 ± 0.418, n=27) compared to control OE (3.6 ± 0.170, n=20) (*p*=0.004). There was no statistically significant difference in mean number of neurites between Aurora Kinase B OE (3.6 ± 0.400, n=26, *p*=1.000) and Aurora Kinase C OE (4.5 ± 0.332, n=29, *p*=0.429) as compared to control OE (3.6 ± 0.170, n=20). E) Sholl analysis shows increase in branching complexity in Aurora Kinase A, B, and C overexpressing cells proximally, with a decrease in branching complexity distally as compared to control OE.

### Overexpression of Nuak Kinases 1 or 2 in Primary Cortical Neurons Suggests Roles for Nuak Kinase 1 in Neurite Initiation and Dendritic Branching

To clarify the role of the different Nuak kinase isoforms overexpression in neurite formation, mouse primary cortical neurons were used to overexpress the Nuak kinases 1 or 2. Following overexpression, neuronal morphology was assessed to analyze the length of neurites likely to become dendrites and neurites likely to become the axon, number of neurites, and arborization (Fig. 5A). Length of neurites likely to become dendrites and neurites likely to become axons was assessed, but no significant differences were found (Fig. 5 B and C). Neurons overexpressing Nuak Kinase 1 (5.5 ± 0.483) had significantly more neurites as compared to the control (3.5 ± 0.312, n=20) (Fig. 5D). Branching was also dramatically increased proximally in neurons overexpressing Nuak Kinase 1 and slightly increased proximally in neurons overexpressing Nuak Kinase 2 (Fig. 5E). Additionally, overexpression of both Nuak kinases resulted in a slight decrease in arborization more distally, with branching that was similar to controls at the most extremely distal lengths (Fig. 5E). Another interesting aspect of morphology is the turning of the neurite likely to be the axon back towards the cell body and axonal swellings in neurons overexpressing Nuak kinase 1 (Fig. 5A, middle panel). Thus, these data suggest that Nuak kinase 1 is involved in neurite initiation and both Nuak kinase 1 and 2 are involved in dendritic branching.

**Figure 5.**
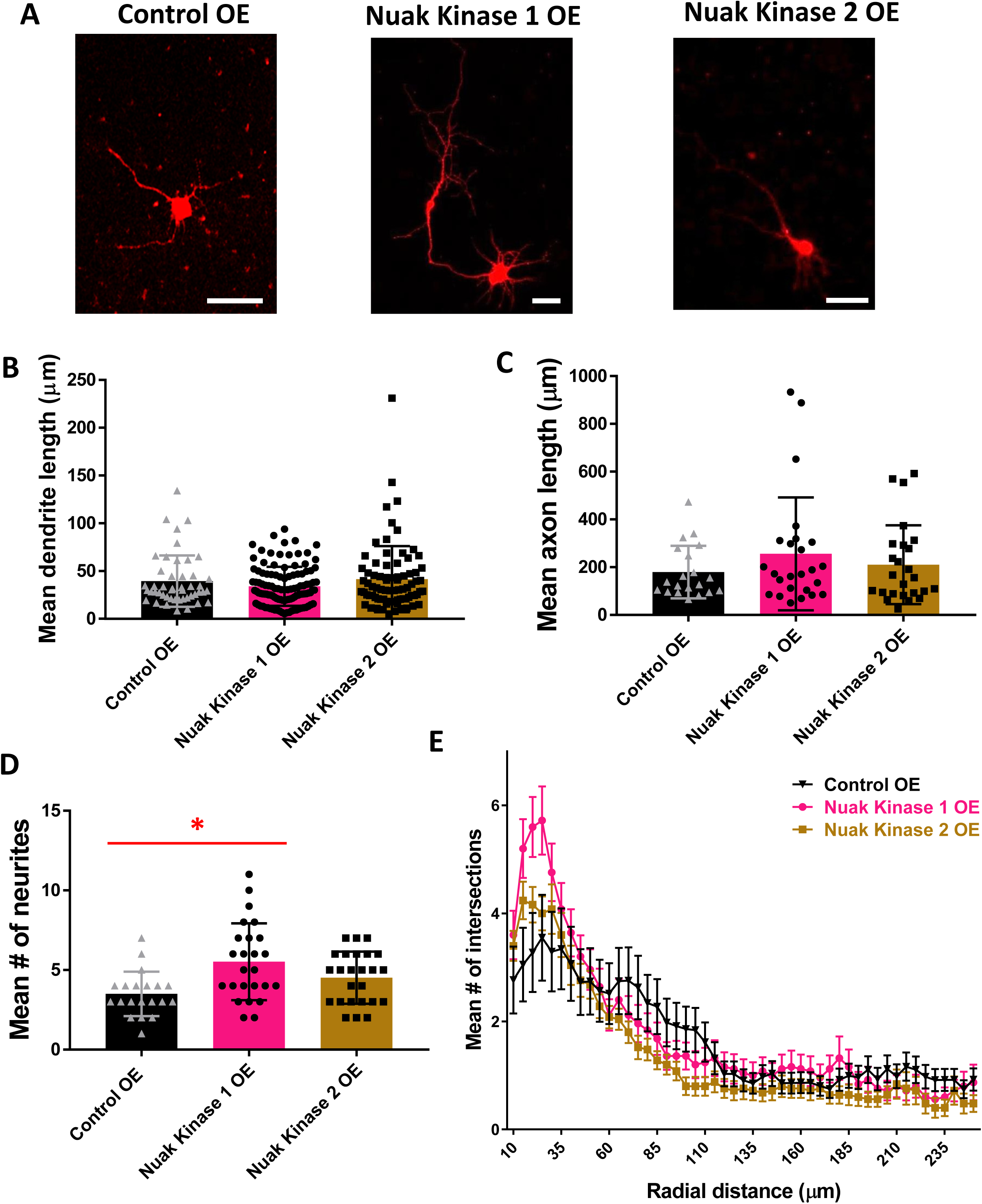
Overexpression of Nuak Kinases 1 and 2 in Primary Cortical Neurons Suggests Roles for Nuak Kinase 1 in Neurite Initiation and Dendritic Branching. A) Primary cortical mouse neurons transfected with mRFP at 48 hours after re-plating, scale bars=50 μm. B) One-way ANOVA determined there was no statistically significant difference in length of neurites likely to become dendrites (F(2,234) = 1.904, *p*=0.151). There was no statistically significant difference between Nuak Kinase 1 OE (34.06 ± 1.94, n=111, *p*=0.742) and Nuak Kinase 2 OE (41.56 ± 3.89, n=79, *p*=1.000) as compared to control OE (39.53 ± 3.92, n=47). C) One-way ANOVA determined there was no statistically significant difference in length of neurites likely to become axons (F(2,67) = 1.010, *p*=0.370). There was no statistically significant difference between Nuak Kinase 1 OE (255.91 ± 47.27, n=33, *p*=0.500) and Nuak Kinase 2 OE (210.43 ± 33.01, n=25, *p*=1.000) as compared to control OE (179.47 ± 24.55, n=20). D) One-way ANOVA determined there was a statistically significant difference (F(2,67) = 6.313, *p*=0.003). Bonferroni post hoc test revealed that mean number of neurites was statistically significantly greater for Nuak Kinase 1 OE (5.5 ± 0.483, n=25, *p*=0.002) as compared to control OE (3.5 ± 0.312, n=20). There was no statistically significant difference in mean number of neurites between Nuak Kinase 2 OE (4.5 ± 0.327, n=25, *p*=0.233) as compared to control OE (3.5 ± 0.312, n=20). E) Sholl analysis shows a dramatic increase in branching complexity in Nuak Kinase 1 overexpressing cells proximally and a slight increase in branching complexity in Nuak Kinase 2 overexpressing cells proximally, with a decrease in branching complexity more distally in both Nuak Kinases as compared to control OE. Branching appears to become similar to controls in both experimental groups at the most distal points.

## Discussion

Our study reports a high-throughput kinase inhibitor screening, which implicates 222 kinase inhibitors as potential therapeutics. There is huge therapeutic potential for manipulation of kinase activity, as nearly 50 kinase inhibitors have already been approved by the FDA for use [10]. Most kinases that were targets of these inhibitors were members of the tyrosine kinase/adaptors, JAK/STAT, membrane transporter/ion channel, and chromatin/epigenetics signaling pathways, highlighting the importance of these pathways in neuritogenesis. We also report roles for Aurora Kinase A in neurite initiation, for Aurora kinases A, B, and C in dendritic branching, and for all Nuak Kinases in neurite initiation and elongation and dendritic branching.

Our screening indicated a role for several pathways that are not traditionally thought to have a role in neurite formation including the tyrosine kinase/adaptors, JAK/STAT, membrane transporter/ion channel, and chromatin/epigenetics signaling pathways pathway in neuritogenesis. Several tyrosine kinases such as anaplastic lymphoma kinase (ALK) have known roles in neurite outgrowth and activated Cdc42-associated tyrosine kinase (ACK1) plays a role in neurite extension and branching [24, 25]. The JAK/STAT pathway is typically thought of as negatively regulating neurite formation and acts through inhibitory proteins such as SOCS2, 3, and 6 [26-28]. Kinases in the membrane transporter/ion channel pathways may regulate ion channels that are important for neurite formation. For example, calcium signaling is known to be an important regulator of neuritogenesis, since the rate of neurite extension is regulated by the frequency of spontaneous calcium transients [29-31]. Kinases in chromatin/epigenetics signaling pathways serve to regulate neurite formation through chromatin remodeling or post-translational modifications of transcription factors [32-34]. Other kinases in these pathways should be investigated more closely to determine if additional regulators of neurite formation may be identified.

Several different phenotypes were observed after treatment with different Aurora and Nuak kinase inhibitors, which acted on different members or combinations of members of Aurora and Nuak kinases. However, it is hard to define the functions of each isoform. In general, our results suggest that the specific combination of kinases inhibited within each family strongly impacts the neurite morphology phenotype seen. The phenotypes seen after inhibition of the Aurora kinases are difficult to interpret, with defects in both neurite initiation and neurite elongation resulting from inhibition of the same kinase(s). Potentially, inhibiting different combinations of Aurora Kinase A, B, and/or C could have caused different compensatory pathways to be activated within the cells. Interestingly, the kinase inhibitor screening in N-2a cells implicated Aurora Kinase A in neurite elongation, while our results in primary cortical neurons show no significant difference in length for neurites likely to become dendrites. This could be due to differences in specific cell types and induction method to extend neurites.

To expand upon our findings from the N-2a cell screening, we used primary cortical neurons and pharmacologically inhibited several individual Aurora and Nuak kinases, as well as multiple different combinations of kinases within each family. Interestingly, inhibition of Aurora Kinase A, which was the only Aurora Kinase to show an overexpression phenotype, resulted in the least severe inhibition phenotype, with no significant difference seen in neurite initiation or elongation and only a mild defect in arborization (Table 4). Combined inhibition of Aurora Kinase A/B may only impact neurite initiation and arborization (Table 4). Inhibition of only Aurora Kinase B may specifically affect neurite elongation and arborization, while combined inhibition of Aurora Kinase B/C has a more severe phenotype compared to inhibition of Aurora Kinase A alone (Table 4). These results suggest that Aurora Kinase B plays a crucial role in regulating neuromorphogenesis, and that Aurora Kinase C also has functions in neuromorphogenesis. Inhibition of either Nuak Kinase 1 or Nuak Kinase 1/2 resulted in severe defects in neurite initiation, neurite elongation, and arborization (Table 4). However, we did not see any significant differences between neurons treated with the Nuak Kinase 1 inhibitor as compared with the Nuak Kinase 1/2 inhibitor. This suggests that Nuak Kinase 1 and 2 may have redundant functions or no functions during neuromorphogenesis, since inhibition of Nuak Kinase 2 in addition to Nuak Kinase 1 does not appear to have any additive effects. These results highlight the crucial need to distinguish effects of kinase targeting drugs on different kinase isoforms and in different cell types. Attention must be paid to isoform specificity and how targeting different combinations of isoforms may lead to redundant or divergent effects.

**Table 4:** Summary of Aurora and Nuak kinases in regulating neuritogenesis and arborization. Summary table of the phenotypes observed in primary cortical neurons when each Aurora and Nuak kinase was either overexpressed or inhibited. X = no significant difference

| **Table 4:** Summary of Aurora and Nuak kinases in regulating neuritogenesis and arborization |  |  |  |  |
| --- | --- | --- | --- | --- |
|  | **Length of**<br>**Neurites likely**<br>**to become**<br>**Dendrites**<br>**(Elongation)** | **Length of**<br>**Neurites likely**<br>**to become**<br>**Axons**<br>**(Elongation)** | **Neurite**<br>**Initiation** | **Arborization** |
| **Aurora A**<br>**overexpression** | X | X | Increased | Increased<br>proximally,<br>decreased<br>distally |
| **Aurora A**<br>**inhibition** | X | X | X | Mild decrease |
| **Aurora B**<br>**overexpression** | X | X | X | Increased<br>proximally,<br>decreased<br>distally |
| **Aurora B**<br>**inhibition** | Decreased | Decreased | X | Decreased |
| **Aurora A/B**<br>**inhibition** | X | X | Decreased | Decreased |
| **Aurora B/C**<br>**inhibition** | Decreased | Decreased | Decreased | Decreased |
| **Aurora A/B/C**<br>**inhibition** | Decreased | Decreased | Decreased | Decreased |
| **Aurora C**<br>**overexpression** | X | X | X | Increased<br>proximally,<br>decreased<br>distally |
| **Nuak 1**<br>**overexpression** | X | X | Increased | Increased<br>proximally,<br>decreased<br>distally |
| **Nuak 1**<br>**inhibition** | Decreased | Decreased | Decreased | Decreased |
| **Nuak 2**<br>**overexpression** | X | X | X | Increased<br>proximally,<br>decreased<br>distally |
| **Nuak 1/2**<br>**inhibition** | Decreased | Decreased | Decreased | Decreased |

The known roles of Aurora kinases A, B, and C during neurite formation, axon regeneration, and mitosis are similar yet distinct. Aurora kinase A has a known role in neurite elongation, and our overexpression results further implicate Aurora kinase A in neurite initiation [13]. Mori et al. found that depletion of Aurora kinase A and any components of the signaling pathway resulted in decreased frequency and speed of microtubule emanation from the microtubule organizing center and into primitive neurites, and subsequent decreased neurite elongation [13]. Microtubule invasion into primitive neurites during neurite initiation is also crucial for neurite stabilization, dictating which lamellipodia and filopodia remain to undergo neurite elongation or retract.

Because Mori et al. performed neurite analysis 48 hours after transfection without re-plating, they likely were only able to observe elongation deficits [13]. Neurite initiation begins immediately upon plating before the transfection has taken full effect, and this experimental timing difference explains the further deficits noted in our Aurora kinase A overexpression experiments which analyzed neurite initiation and elongation 48 hours after re-plating after the transfection has reached full efficiency. Elongation results from Aurora kinase A inhibition in primary cortical neurons is in agreement with previous literature in dorsal root ganglion cells, although we show a trend towards decreased neurite length that does not reach significance. We observed increased neurite initiation but did not observe changes to neurite elongation upon Aurora kinase A overexpression. Overexpression is not a measure of protein function but can be useful to test the effects of manipulation of certain targets. This suggests that although Aurora kinase A has important developmental roles in neurite elongation, its activation may not be useful to therapeutically target axon elongation of a damaged axon. Instead, it could be applied to axon re-initiation in cases where the axon must re-emerge from the soma.

Aurora kinase B has a known role in axon outgrowth and regeneration in zebrafish [14]. Gwee et al. found that pharmacological inhibition of Aurora kinase B prevented both axonal outgrowth and regeneration in spinal motor neurons, and that overexpression of Aurora kinase B led to increased axon elongation [14]. Our kinase inhibitor results are in agreement with these results, and also revealed further deficits that were not specific to the axon but to all neurites. We found that Aurora kinase B inhibition results in decreased dendrite and axon length in mouse primary cortical neurons. It is possible that cell and species type specific activity of Aurora kinase B contributed to our additional results. Taken together, our study and others have provided strong evidence that Aurora kinase B contributes to neurite elongation in post-mitotic cells. Surprisingly, overexpression of Aurora kinase B had no significant effects on cortical neuron morphology. Aurora kinase B’s canonical role in mitosis maintains chromosome alignment and segregation fidelity [43]. One of the many ways Aurora kinase B promotes these mechanisms is through phosphorylation and activation of microtubule depolymerizer, Kif2A, which was recently shown to contribute to neuronal morphology [44]. It has been shown that in cases where Aurora kinase B is altered during development, endogenous activity of Aurora kinase C can rescue these deficits [45]. It is not unlikely that a compensatory mechanism of this nature could also take place during overexpression of either kinase.

Aurora kinase C has overlapping canonical roles with Aurora kinase B [43]. We are the first to analyze the role of Aurora kinase C in post-mitotic cells. We discovered that Aurora kinase C contributes to dendritic branching, implicating its inhibition as a therapeutically valuable target to trim back aberrant neurite branching following seizures. Because none of our kinase inhibitors were selective for only Aurora kinase C, it is hard to interpret whether Aurora kinase C is important for neurite formation as the results seen in our inhibitor assay could be due to combined inhibition of Aurora kinase B. Since Aurora B and C have similar roles during mitosis, with one of their major functions being to organize microtubules, we may speculate that Aurora B and C may also function similarly to each other during neurite formation. This may explain why combined inhibition of Aurora B and C leads to defects. Additionally, Aurora A functions differently during mitosis, with one of its main roles being to regulate microtubule assembly and polymerization. This could explain why we only see an overexpression phenotype when Aurora A is overexpressed, since increased microtubule polymerization could increase neurite outgrowth, while increasing microtubule organizing regulators, which may be occurring when Aurora B and C are overexpressed, may not impact neurite formation [46]. Furthermore, Aurora B and C have known compensatory roles for each other when one is disrupted. Our results suggest differential roles for kinases in the Aurora family. This information could be useful in designing therapeutics for targeting either axon or dendrite degeneration or regeneration. For example, aberrant neurite formation may occur following seizure, resulting in the formation of inappropriate synaptic contacts [47]. A therapeutic may be designed to inhibit neurite initiation, elongation, and/or branching after seizure. Additionally, alterations in dendritic branching are related to Rett Syndrome and Fragile-X[48]. Developing a therapeutic to modulate the extent of branching may improve clinical phenotypes. After identifying targets with potential for further investigation, we might next investigate which structural domains are relevant for affecting neuronal morphology. While the Aurora kinases are structurally similar, there are notable differences. Aurora Kinase B and Aurora Kinase C share 75% homology of entire sequence and kinase domain, while Aurora Kinase A and Aurora Kinase C share 60% homology of total sequence and kinase domain [49]. Future studies may identify which domains are involved in the regulation of axon and dendrite length by mutating several regions of each kinase that have unique homology to determine what effect each domain has on regulating neuromorphogenesis.

Although the overexpression of the kinases of interest is not a direct functional analysis, results from overexpression can suggest the functions of the protein and/or the utility of activating or overexpressing that protein. Our overexpression experiments indicate Aurora Kinase A and Nuak Kinase 1 are critical factors during neurite initiation and dendritic branching. However, to define the functions of each isoforms in more detail, a specific loss-of-function analysis such as shRNA knockdown should be performed.

Although previous literature first reported a role for the Aurora kinases A and B and Nuak 1, our overexpression phenotypes confirm these results as well as implicate the Aurora and Nuak kinases in additional aspects of neuromorphogenesis. Aurora Kinase A has a known role in neurite elongation and Aurora Kinase B has a known role in regulating axon length [12-14]. We show here that Aurora Kinase A is also involved in neurite initiation and that the Aurora kinases A, B, and C are all involved in dendritic branching. Previously, Nuak Kinase 1 has been reported to regulate axonal branching [16]. We describe roles for Nuak Kinase 1 in neurite initiation and dendritic branching as well. Our results further emphasize the importance of the Aurora and Nuak kinases in multiple steps of neurite formation and dendritic branching.

In conclusion, our results support the previous observations reported from other groups, and novel roles for the Aurora and Nuak kinases in regulating neurite initiation and dendritic branching have been clarified. Inhibition and overexpression phenotypes also reveal differences within these kinase families, suggesting that each class of kinase plays a very specific role in establishing axons and dendrites. Additionally, the high-throughput kinase inhibitor screening we conducted has the potential to call attention to new targets of investigation in the search for better therapeutics to treat patients suffering from aberrant axon sprouting or failure to regenerate injured axons.

## Methods

### Mice

All animal experiments were performed under protocols approved by the Drexel University Animal Care and Use Committees and following the guidelines provided by the US National Institutes of Health. C57BL/6 mice were maintained in our animal facility and used for setting up mating to obtain the time-pregnant mice. Embryonic day (E) 0.5 was defined as the morning of the day the vaginal plug appeared.

### Kinase Inhibitor Library

DiscoveryProbeTM kinase inhibitor library (Cat # L1024) was purchased from APExBIO (Boston, MA). The library is a collection of 493 kinase inhibitors, mainly including inhibitors, but also some activators. Original 10mM in DMSO stocks were stored at −80°C, and 1mM working solutions were created by diluting the stock solution in DMSO and also storing at −80°C until use. The full list of inhibitors is supplied as a supplemental table (Suppl. Table 2).

### Kinase Inhibitor Screening

Neuro-2a (N-2a) mouse neuroblastoma cells were plated in a 96 well plate that was coated with poly-L-lysine (PLL). N-2a cells were cultured for 1 hour in DMEM (+) media containing kinase inhibitors from the DiscoveryProbeTM Kinase Inhibitor Library (APExBIO). After 1 hour, the media was changed to DMEM (-) with inhibitors to induce neurite outgrowth. Each of the 493 kinase inhibitors investigated were tested at concentrations of 0.1, 1, and 10 μM to optimize the dosage, since there was a range of doses at which the kinase inhibitors could be administered before a toxic level was reached and cell death was observed. After 3 hours cells were fixed. Three blinded observers used a scoring system to rank the extent of neurite formation in each condition by assigning each a score from 0-8, with 0 indicating a total absence of neurites and 8 indicating long neurites on almost all cells (Table 1).

### Calculation of Scores for Kinase Inhibitor Phenotypes

Scores from three blind observers were averaged. The average control score for each individual 96 well plate differed slightly, since some individual plates had slightly longer or shorter neurites in general. For the kinase inhibitor screening experiment, we used eighteen 96 well plates, and there were 8 control wells on each 96 well plate. The mean control phenotype for all 18 of the 96 well plates was 5.04. The average of all controls (5.04) was subtracted from the average score for each plate to determine the amount that the mean scores would be adjusted by in order to normalize the scores. This value was then subtracted from the mean experimental scores for each kinase inhibitor to normalize each score. The following calculation was done to control for the slight differences in neurite length between each individual plate: ((Observer 1 Score + Observer 2 Score + Observer 3 Score)/3) – (Plate Specific Mean Control Score – 5.04) = Normalized Score.

### Plasmids

pCAG-mRFP was a gift from Joseph Loturco (Addgene plasmid # 28311; http://n2t.net/addgene:28311; RRID:Addgene_28311). pCAG-YFP was a gift from Connie Cepko (Addgene plasmid # 11180; http://n2t.net/addgene:11180; RRID:Addgene_11180). CDNA expression plasmids for Aurora kinases A, B, and C and Nuak kinases 1 and 2 were purchased from OriGene Technologies, Inc., (Rockville, MD). N-terminal Myc/DDK-tagged cDNAs in pCMV6-Entry vector were used. All plasmids were purified using NucleoBond Xtra purification kit (MACHEREY-NAGEL Inc., Bethlehem, PA).

### Primary Mouse Cortical Neuronal Culture and Overexpression of Aurora and Nuak Kinases

Primary mouse cortical neurons were harvested from embryonic mice at E15.5, prior to gliogenesis at E17.5, to allow for the exclusive culture of neurons [61]. Neurons were plated in wells coated with PLL and laminin. Co-transfection of mRFP + Aurora kinases A, B, and C, and mRFP + Nuak 1 and 2 kinases was performed via chemical transfection using TransIT-X2 (Mirus Bio, Madison, WI) 48 hours following plating. Neurons were re-plated on coverslips coated in PLL and laminin and fixed using 4% PFA after an additional 48 hours. Re-plating allowed neurite formation to occur after the transfection had taken full effect.

### Treatment of Primary Mouse Cortical Neurons with Aurora and Nuak Kinase Inhibitors

Primary mouse cortical neurons were harvested and cultured as described above. Neurons were transfected with YFP via nucleofection. Neurons were treated with Neurobasal media containing 5 μM of kinase inhibitor from the DiscoveryProbeTM Kinase Inhibitor Library (APExBIO). The following inhibitors were used: MK-5108 (VX-689) – highly selective inhibitor of Aurora Kinase A, Hesperadin – Aurora Kinase B inhibitor, TAK-901 – Aurora A/B inhibitor, GSK1070916 – Aurora B/C inhibitor, SNS-314 Mesylate – potent and selective inhibitor of Aurora A/B/C kinases, HTH-01-015 – highly specific and selective Nuak 1 inhibitor, WZ4003 – potent and selective Nuak 1/2 inhibitor. Controls were treated with DMSO, as the inhibitors were diluted in DMSO. Neurons were fixed 48 hours after re-plating using 4% PFA.

### Analysis of Neuronal Morphology

ImageJ software measuring features were used to analyze neuronal morphology of primary cortical neurons. Maximum projection images from z-projection photos produced from z-stack data collected by confocal microscope (Leica SP2 and SP8) were used for analysis. Sholl analysis was performed using the Sholl Analysis Plugin (Gosh Lab, UCSD) for ImageJ following the developer instructions. The longest neurite was considered the neurite most likely to become the axon, while the remaining neurites were considered neurites likely to become dendrites.

## Statistical Analysis

Quantitative data were subjected to statistical analysis using SPSS (IBM Analytics). The data were analyzed by one-way ANOVA with Bonferroni when appropriate. Values represented as mean ± S.E.M. Results were deemed statistically significant if the *p* value was <0.05.

## Acknowledgement

This work has been supported by a research grant from the NINDS (NS096098).

## Author contributions

SMB performed some experiments and data analysis and wrote a draft of the manuscript. SAB and XL performed some experiments and data analysis. KT edited the manuscript and finalized it.

## Supplemental Tables

**Supplemental Table 1. Complete Screening Data for 493 Kinase Inhibitors**

A) Raw scores for each kinase inhibitor at concentrations of 0.1, 1, and 10 μM from three blind observers. Scores were averaged and normalized as described in Methods. B) Raw data classifying each inhibitor concentration as resulting in either no phenotype, neurite initiation defect, neurite elongation defect, or negative regulation. C) Phenotype summary of dose-dependent inhibitors organized by signaling pathway.

**Supplemental Table 2. Kinase Inhibitor Library Information**

A) Chemical data for all kinase inhibitors. B) Table showing the number of inhibitors in the library that target each kinase.

